# Proline-rich Extensin-like Receptor Kinases PERK5 and PERK12 are involved in Pollen Tube Growth

**DOI:** 10.1101/2021.01.12.425807

**Authors:** Cecilia Borassi, Ana R. Sede, Martin A. Mecchia, Silvina Mangano, Eliana Marzol, Silvina P. Denita-Juarez, Juan D. Salgado Salter, Silvia M. Velasquez, Jorge P. Muschietti, José M. Estevez

## Abstract

**Background:** Cell wall integrity plays an essential role during polarized cell growth typical of pollen tubes and root hairs. <u>P</u>roline-rich <u>E</u>xtensin-like <u>R</u>eceptor <u>K</u>inases (PERK) belong to the hydroxyproline-rich glycoprotein (HRGP) superfamily of cell surface glycoproteins.

**Results:** Here, we identified two *PERKs* from *Arabidopsis thaliana*, PERK5 and PERK12 highly expressed in mature pollen. Pollen tube growth was impaired in the single and double *perk5-1 perk12-1* loss of function mutants, with a moderate impact on seed production. When the segregation of self- and reciprocal-crosses of the *perk5-1*, *perk5-2* and *perk12-1* single mutants, and reciprocal-crosses of the *perk5-1 perk12-1* double mutant were carried out, a male gametophytic defect was found, indicating that *perk5-1* and *perk12-1* mutants carry defective pollen tubes, resulting in deficient pollen transmission. Furthermore, double *perk5-1 perk12-1* mutants show excessive accumulation of pectins and cellulose at the cell wall pollen of the tube tip. In addition, an upregulation of cytoplasmic ROS levels were detected by using 2,7-dichlorofluorescein diacetate probe (H_2_DCF-DA), and in agreement, similar results were obtained with HyPer, a genetically encoded YFP-based radiometric sensor, which is used to follow the production of hydrogen peroxide (H_2_O_2_). Single and double *perk5-1 perk12-1* mutants show higher levels of cytoplasmic H_2_O_2_ in their pollen tube tips.

**Conclusions:** Taken together, our results suggest that PERK5 and PERK12 are necessary for proper pollen tube growth highlighting their role on cell wall assembly and ROS homeostasis.

## Background

Pollen grains are required for sexual reproduction which plays a vital role in seed formation and, therefore, in the productivity of crops. The main pollen function is to generate and transport sperm cells to the ovules (Borg et al. 2009). In compatible interactions, the pollen grain germinates on the stigma generating a pollen tube that grows through the transmitting tissue of the style, until reaching the ovule where double fertilization occurs (Crawford and Yanofsky, 2008; Dumas and Rogowsky, 2008). Pollen tube growth is a polarized cell expansion process, and requires an oscillatory positive feedback loop of calcium ions (Ca^2+^), Reactive Oxygen Species (ROS) and pH. Polar-growth relies on the stretching of the existing primary cell wall in the apical zone accompanied by secretion of new cell wall materials (Altartouri & Geitmann 2015). Overall, ROS, Ca^2+^ and pH oscillations are coupled to a transient cell wall loosening to allow a turgor-driven localized cell expansion (Braidwood et al, 2014; Wolf & Höfte 2014). Pollen tube cell walls are enriched in pectins, glycoproteins and xyloglucans/cellulose and deficiencies in any of these polymers at the cell wall inhibit polar cell elongation, indicating that they operate in a coordinately manner to modulate tip growth (Dardelle et al, 2010; Gu and Nielsen, 2013; Sede et al, 2018; Wang et al. 2018).

Polar growth must be tightly regulated to allow the maintenance of cell wall integrity (CWI) and coordination during plant development. Plant cells have developed a rich diversity of complex proteins with diverse extracellular domains connected to intracellular domains to convey environmental and cell wall signals within the cell (Ringli, 2010; Borassi et al. 2015). In *A. thaliana* Receptor-like kinases (RLKs) comprise ~600 members mostly localized in the plasma membrane (PM) mediating extracellular signals to the cytoplasm and nucleus (Shi et al, 2004; Wolf et al, 2012; Muschietti & Wengier 2018). At least four RLK subfamilies have been implicated in sensing CWI during cell expansion: Wall-Associated Kinases (WAKs), Lectin Receptor Kinases (LecRKs), *Catharanthus roseus* Receptor-Like Kinase1-Like proteins (CrRLK1Ls), and Proline-rich, Extensin-like Receptor Kinases (PERKs). PERK proteins consists of an extracellular domain, a typical transmembrane and an intracellular kinase domain where the kinase activity resides. The extracellular domain is rich in contiguous prolines, some of them are part of a classical Extensin (EXT)-motif with SerPro_(3–5)_ repeats but lack an adjacent YXY for Tyr-mediated protein crosslinking. In *Arabidopsis thaliana*, the PERK family contains 15 related-members (*AtPERK1-15*) (Silva and Goring, 2002; Nakhamchik et al. 2004). Nine of them are highly expressed in expanding root hairs (e.g. *PERK8* and *PERK13*) and pollen tubes (e.g. *PERK3-7,11-12*) suggesting an specialized and unique role of PERKs in polar based growth (Borassi et al. 2015; Chen et al. 2020; Li et al. 2020). The presence of EXT-domains in the apoplastic side would suggest PERKs as putative sensors of the EXT-pectin glyco-network (Cannon et al. 2008; Marzol et al. 2018; Herger et al. 2020) as it was demonstrated for WAKs in regards to pectins (Kohorn, 2015). Although PERKs have been connected with polarized cell expansion in root hairs (Won et al., 2009; Hwang et al., 2016) there is still no evidence of their function during polarized growth of pollen tubes. *PERK13* (*RHS10* for Root Hair Specific 10) is specifically expressed in root hairs and modulates the duration of root hair polar-growth and thus its length (Won et al., 2009; Hwang et al., 2016). *rhs10* mutant produces longer root hairs as well as higher ROS accumulation than the WT, although the molecular mechanism is still unclear (Hwang et al., 2016). PERK4 is involved in Ca^2+^ signaling and abscisic acid response in root tip growth (Bai et al. 2009a, b). Based on interaction studies and mutant phenotypes, PERK8/9/10 kinase domains together with the AGC VIII kinase (AGC1-9) and the closely related kinesin-like calmodulin-binding protein (KCBP)-interacting protein kinase (KIPK) may function together to trigger a signaling response during root growth (Humphrey et al., 2015). Other members of PERK family have been related to apical dominance and root elongation in *Arabidopsis* (Hwang *et al*., 2010; Bai *et al*., 2009; Humphrey et al., 2015), suggesting that PERKs are involved in the regulation of several plant growth and developmental processes. In this work, we show that pollen tubes require both PERK5 and PERK12 for proper pollen tube polar growth linked to the cell wall polysaccharide assembly, most probably pectins and cellulose, as well as to ROS homeostasis. Double *perk5 perk12* knock out mutant produce shorter pollen tubes probably due to alterations in cell wall polysaccharide composition and ROS levels that affect ovule fertilization. These findings provide an insight into the biological function of PERKs during pollen tube growth and fertilization process.

## Results

Based on pollen transcriptomic studies, several PERK genes are expressed in mature pollen (**Supplementary Figure 1A**). *PERK3* (At3g24540), *PERK4* (At2g18470), *PERK5* (At3g18810), *PERK6* (At4g34440), *PERK7* (At1g49270), *PERK11* (At1g10620) and *PERK12* (At1g23540) are expressed late during pollen development (Honys and Twell, 2004) suggesting a role during pollen tube growth and /or pollen-pistil interactions. In order to confirm their expression in the male gametophyte, the GUS reporter gene was expressed under the control of the respective pollen PERK promoters. As shown in **Figure 1** and **Supplementary Figure 2**, GUS activity of all pollen PERK genes (*PERK3, PERK4, PERK5, PERK6, PERK7, PERK11* and *PERK12*) was found in pollen grains and pollen tubes. However, the GUS signal in pollen tubes could be due to its expression in mature pollen. A GUS signal was only observed when WT or p*PERK5::GUS* pistils were pollinated with p*PERK5::GUS* pollen, but not with WT pollen, demonstrating that *PERK5* is expressed in pollen and not in the style-transmitting tract (**Figure 1B**). No GUS activity was detected in *PERK3, PERK4, PERK5, PERK6, PERK7, PERK11* and *PERK12* 10-d-old transgenic seedlings (**Figure 1** and **Supplementary Figure 2**). All these results suggest that *PERK3, PERK4, PERK5, PERK6, PERK7, PERK11* and *PERK12* are expressed in mature pollen, and possibly in pollen tubes. To characterize the function of the different pollen PERKs, homozygous Arabidopsis lines for T-DNA insertions were isolated. As shown in **Figure 2A** and **Supplementary Figure 1B**, two insertion alleles were selected for *PERK5*, and for the rest of the *PERK* genes, one insertion allele was chosen within the corresponding exons. RT-PCR analysis revealed that *perk3-1, perk4-1, perk5-1, perk5-2, perk6-1, perk7-1, perk11-1*, and *perk12*-mutants do not show expression of the corresponding disrupted genes (**Figure 2B and Supplementary Figure 1C**).

**Figure 1.**
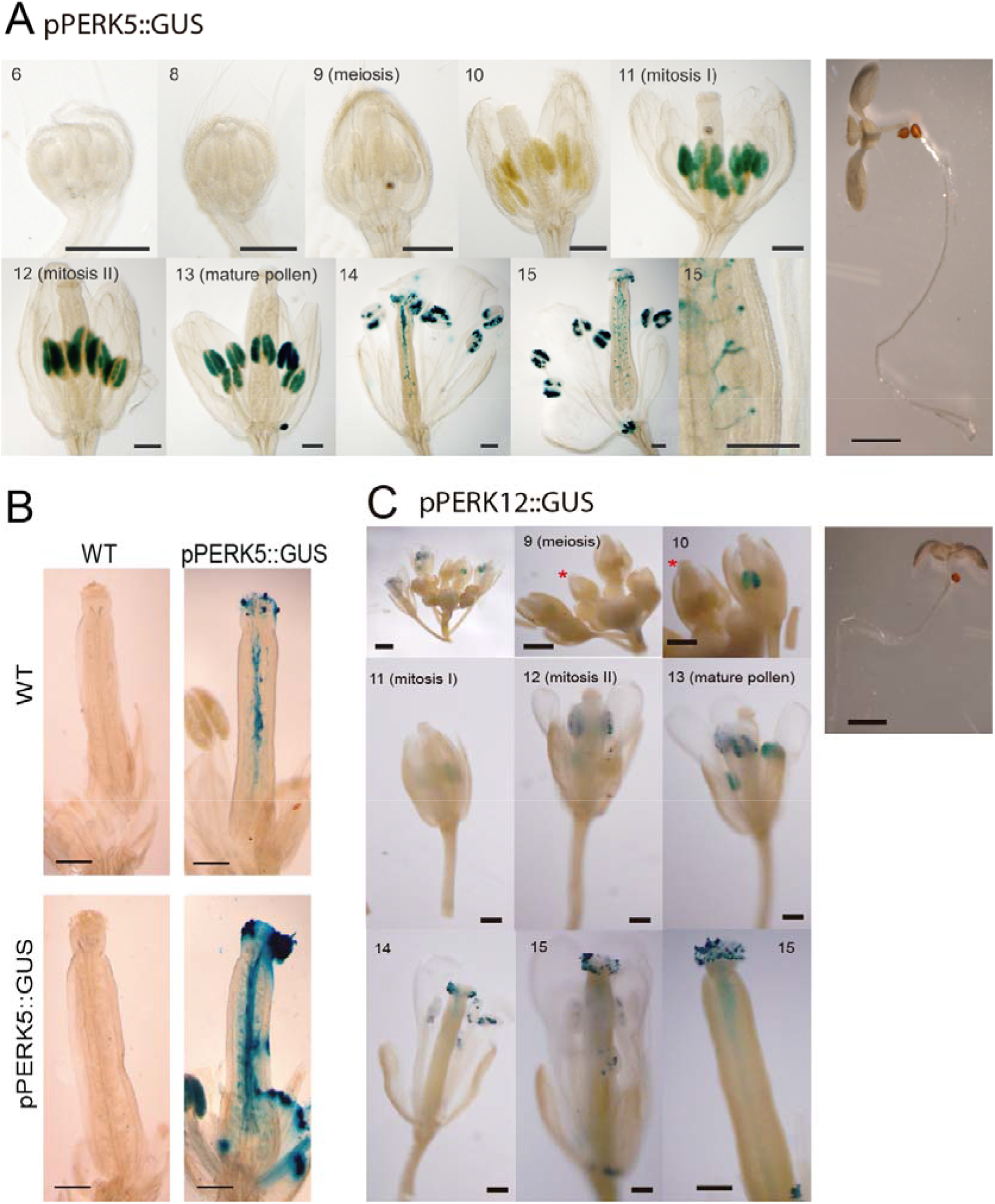
PERK5 and PERK12 are expressed during pollen development. (**A**) Analysis of GUS activity in transgenic plants carrying the construction *pPERK5::GUS*. GUS activity observed in pollen grains within the anther (from developing stages 11 to 15) and in germinated pollen grains on the stigma. Inset on stage 15 shows a pollen tube that reached the ovule. Right panels visualize a seedling in which GUS activity was not detected. Scale bar: 100 μm. (**B**) WT and *pPERK5::GUS* transgenic plants pollinated either with WT or *pPERK5::GUS* pollen grains. GUS staining was not observed in pollen grains or pollen tubes from WT plants. (**C**) GUS activity in *pPERK12::GUS* transgenic plants was examined before (stages 9-12) and after (stages 13-15) anthesis. GUS activity was visualized from stage 11 (mitosis I) to stage 15. Red asterisks indicate the flower that corresponds to the stage displayed in the image. Right panels visualize a seedling in which GUS activity was not detected. Scale bar= 100 μm.

**Figure 2.**
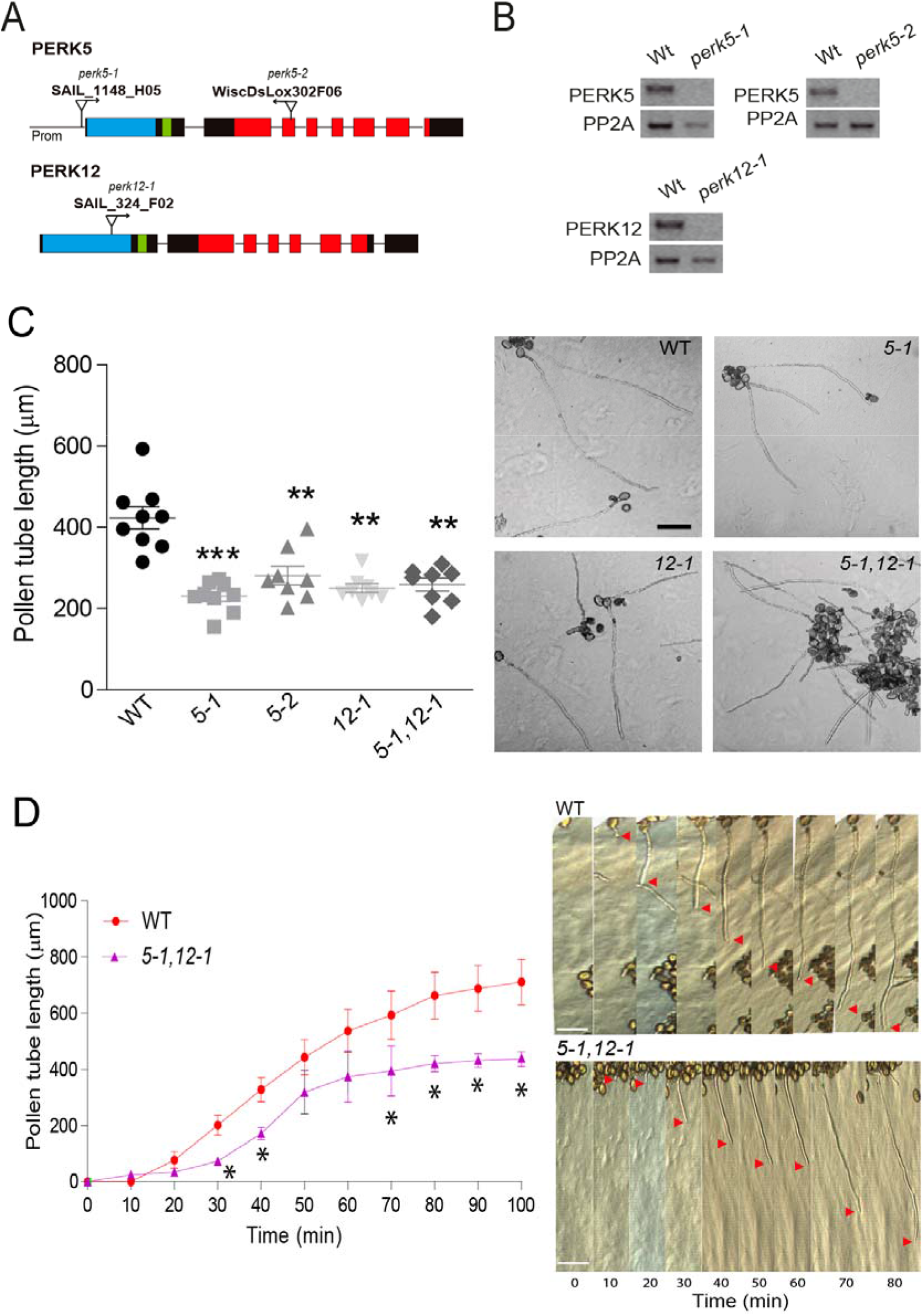
PERK5 and PERK12 are involved in pollen tube *in vitro* growth. (**A**) Schematic representation of *PERK5* and *PERK12* genes showing the extracellular domains (EXT)-motif with SerPrO_(3–5)_ repeats (light blue), transmembrane (green) and kinase domains (red) coding regions. Introns (thin lines), exons (rectangles), and positions of T-DNA insertions are also indicated. Bars = 100 bp. **(B)** RT-PCR of *perk* mutant lines. RNA was extracted from pollen tubes germinated in vitro. PP2A was used as control. (**C**) Pollen tube length of single mutants *perk5-1, perk5-2* and *perk12-1* and double mutant *perk5-1 perk12-1* after 3 h of germination *in vitro*. Data are shown as the mean ± SEM (n=11). Asterisks represent significant differences from WT according to one-way ANOVA test (**) p<0.05, (***) p<0.01. On the right, representative images of the quantification shown in C. Scale bar= 100 mm. (**D**) Kinetics of pollen tubes grown *in vitro* for 100 min. Pollen tube length is reported for each time point as the average of 20 pollen tubes. Data are shown as mean ± SEM and the asterisks represent significant differences from the WT according to Student’s test p <0.05. On the right, representative images of the analysis of pollen tube growth kinetics, for the WT and the double mutant *perk5-1 perk12-1*. Red arrows indicate pollen tube tip. Scale = 100 μm.

To examine whether pollen PERKs are necessary for normal pollen germination and pollen tube growth, mutant pollen was germinated *in vitro* for 3h. **Figure 2C** and **Supplementary Figure 3A** show that only *perk5-1, perk5-2* and *perk12-1* single mutants had shorter pollen tubes when compared to WT pollen. Double mutant analysis showed that only the double mutant *perk5-2 perk12-1*, but not *perk4-1 perk7-1, perk6-1 perk7-1, perk6-1 perk11-1* and *perk7-1 perk11-1*, displayed significant differences in *in vitro* pollen tube growth compared to WT pollen **(Figure 2C and Supplementary Figure 3B)**, suggesting that there is a high degree of functional redundancy within the pollen PERK family. Since pollen tube length of the double mutant *perk5-1 perk12-1* resembled that of both single mutants, *PERK5* and *PERK12* might indeed act redundantly in regulating pollen tube growth. Based on all these results, we selected the *perk5-1, perk5-2* and *perk12-1* single mutants and the *perk5-1 perk12-1* double mutant for deeper analysis. Kinetic analysis of pollen tube growth indicates that *perk5-1 perk12-1* double mutant rate (*perk5-1 perk12-1_vel_* = 143.1 μm/h) was lower than WT (WT_vel_ = 325.7 μm/h) **(Figure 2D)** highlighting a role for *PERK5* and *PERK12* in pollen tube growth.

When the segregation of self-crosses of single mutants *perk5-1, perk5-2*, and *perk12-1* heterozygous plants was analyzed, a statistically significant deviation from the expected 1:2:1 segregation ratio was observed, indicating a gametophytic defect (**Table 1**). Reciprocal crosses were made to analyze whether the reduced transmission was caused by a defect in the male or female gametophyte. **Table 1** shows that impaired segregation was observed only when mutant pollen was used while it was normal through the female gametophyte. All these results suggest that *perk5-1* and *perk12-1* single mutants carry defective pollen tubes, resulting in a deficient pollen transmission.

**Table 1.**
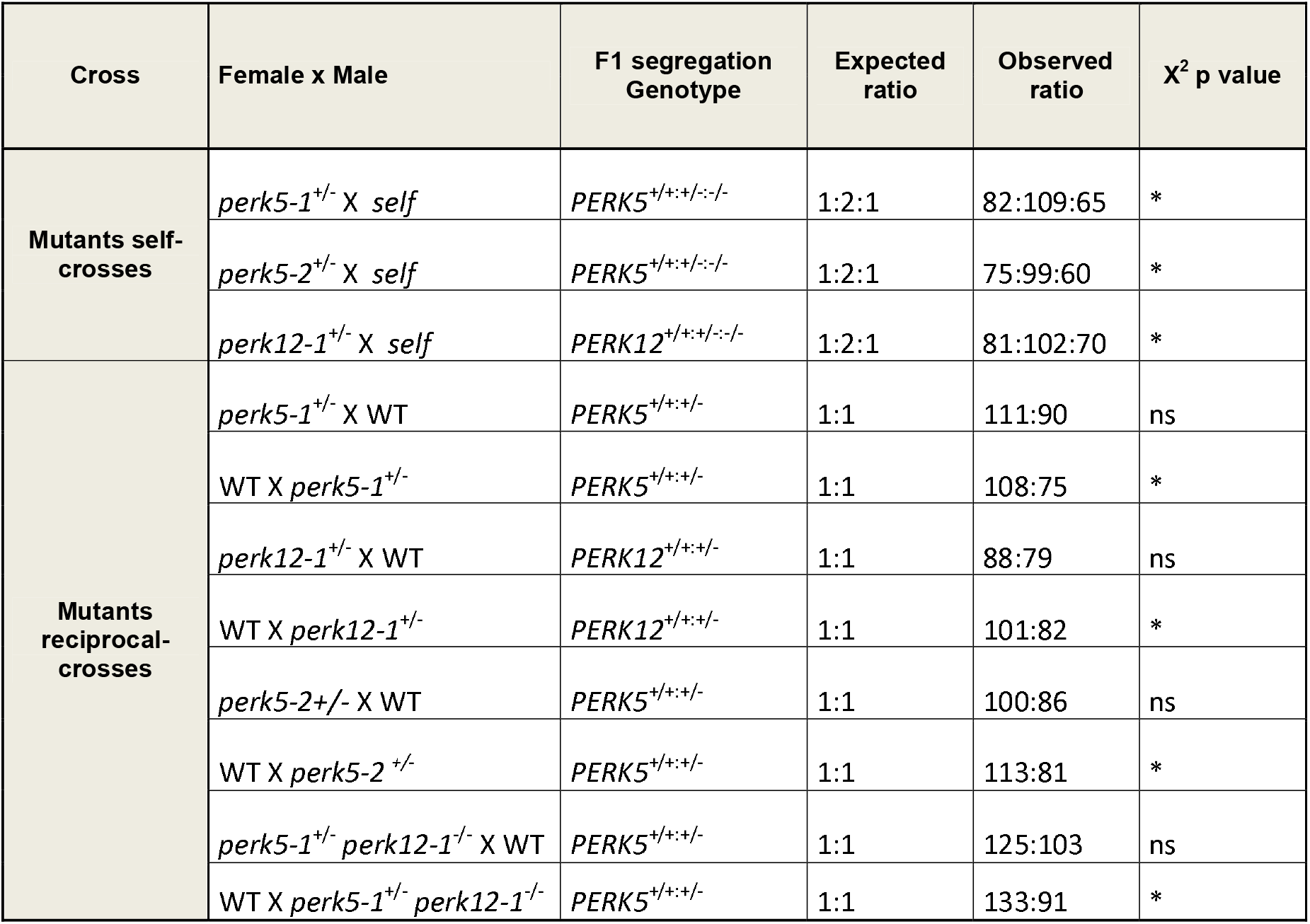
Segregation of self- and reciprocal-crosses of the *perk5-1, perk5-2* and *perk12*-1 single mutants, and reciprocal-crosses of the *perk5-1 perk12-1* double mutant. Differences were evaluated with the Chi-square test (χ2) *p<0,05. ns: no significant differences.

To assess whether the observed reduction in pollen transmission of mutants was due to a deficiency in pollen tube growth, semi *in vivo* and *in vivo* pollen tube growth assays were performed. For the semi *in vivo* analysis, WT or *perk5-1* and *perk12-1*single mutants and double mutant *perk5-1 perk12-1* pistils were hand pollinated, cut at the upper section and placed in semi-solid pollen germination medium waiting for pollen tubes to emerge from the cut style (Palanivelu & Preuss, 2006). No differences in pollen tube growth were observed when WT pollen was used to pollinate either WT or single and double mutant pistils **(Figure 3A)**. In contrast, when mutant pollen was used in WT or mutant pistils, statistically significant shorter pollen tubes emerged from the cut style, demonstrating that pollen tube growth is defective in the single *perk5-1* and *perk12-1* and double *perk5-1 perk12-1* mutants **(Figure 3A)**. When *in vivo* pollen tube growth was analyzed 12 hs after hand pollination, we observed that mutant *perk5-1, perk5-2*, perk12-1 and *perk5-1 perk12-1* pollen tubes were significantly shorter than the WT. **(Figure 3B)**. Analysis of the seed set in all mature siliques of homozygous self-cross mutant plants **(Figure 3C)**, revealed that single mutant *perk5-1, perk5-2* and *perk12-1* and double mutant *perk5-1 perk12-1* plants produce slightly less seeds per silique compared to WT plants, again indicating functional redundancy within the pollen PERK family. Taken together, these results suggest that the fertility defects observed in single and double PERK mutants are exclusively due to deficiencies in the male gametophyte.

**Figure 3.**
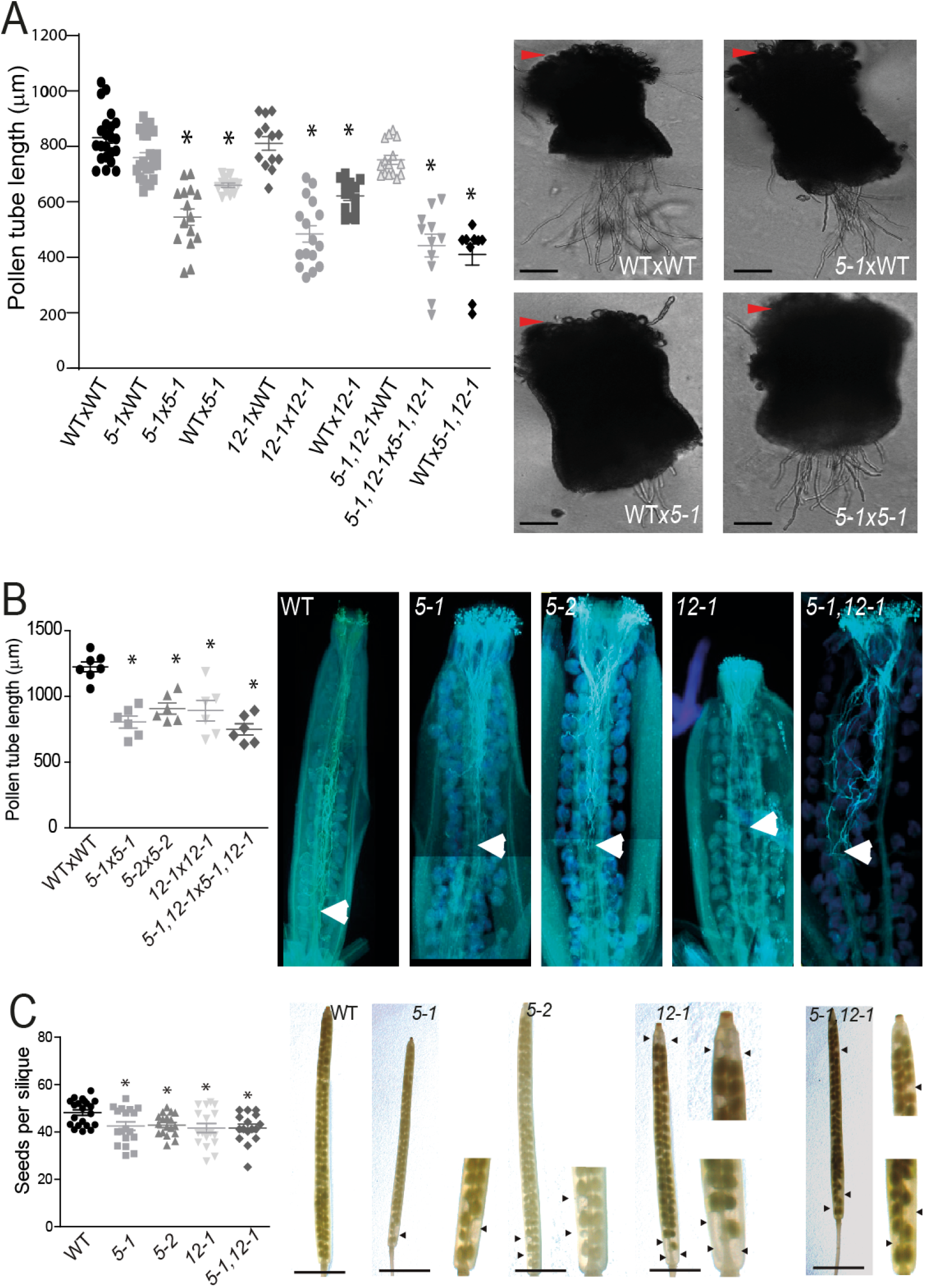
PERK5 and PERK12 are involved in pollen tube growth and their absence impact on the seeds set. (**A**) Quantification of pollen tube length in a *semi-in vivo*assay. Styles were observed 3 h after hand-pollinated. Data are shown as mean ± SEM and the asterisks represent significant differences from the WT according to one-way ANOVA test p <0.05. On the right, representative images of WT and *perk5-1* pollen tubes growing through WT or *perk5-1* mutant right pistils in a *semi-in vivo* assay. Arrowheads indicate pollen grains on the stigma. Scale bar= 100 mm. (**B**) Quantification of pollen tube length in *in vivo* assays using self-pollinated pistils. On the right, representative images of an *in vivo* aniline blue staining of pollen tubes grown for 12h. Arrowheads indicate pollen tube tips inside the pistil. Scale bar= 500 mm. (**C**) Number of seeds per silique in WT, single and double *perk* mutant plants. Data are shown as the mean ± SEM of ≥ 15 plants per genotype (10 siliques were analyzed for each plant). Asterisks indicate significant differences compared to the WT according to one-way ANOVA test with p <0.05. On the right, representative images of the quantification. Scale bar= 500 μm.

PERKs proteins contain extensin (EXT)-motifs in their extracellular domains suggesting a role in sensing changes of cell wall composition. Based on this hypothesis, the pectin and cellulose abundance of WT, single mutant *perk5-1, perk5-2* and *perk12-1* and double mutant *perk5-1 perk12-1* pollen tubes were quantified **(Figure 4 A-B)**. Propidium iodide (PI) which stains pectin, mostly as non-esterified homogalacturonans, and Pontamine Fast Scarlet 4B (S4B) for cellulose detection were used. The PI and S4B signals were quantified along the perimeter of the pollen tubes from the tip to the subapical area **(Figures 4A-B)**. All these *perk* mutants show a significant increase in pectin and cellulose deposition at the cell wall of pollen tubes when compared to the WT. These results suggest that the decrease in growth rate previously observed **(Figure 2D)** might be due to an increase in cell wall stiffness in *perk* mutant pollen tubes.

**Figure 4.**
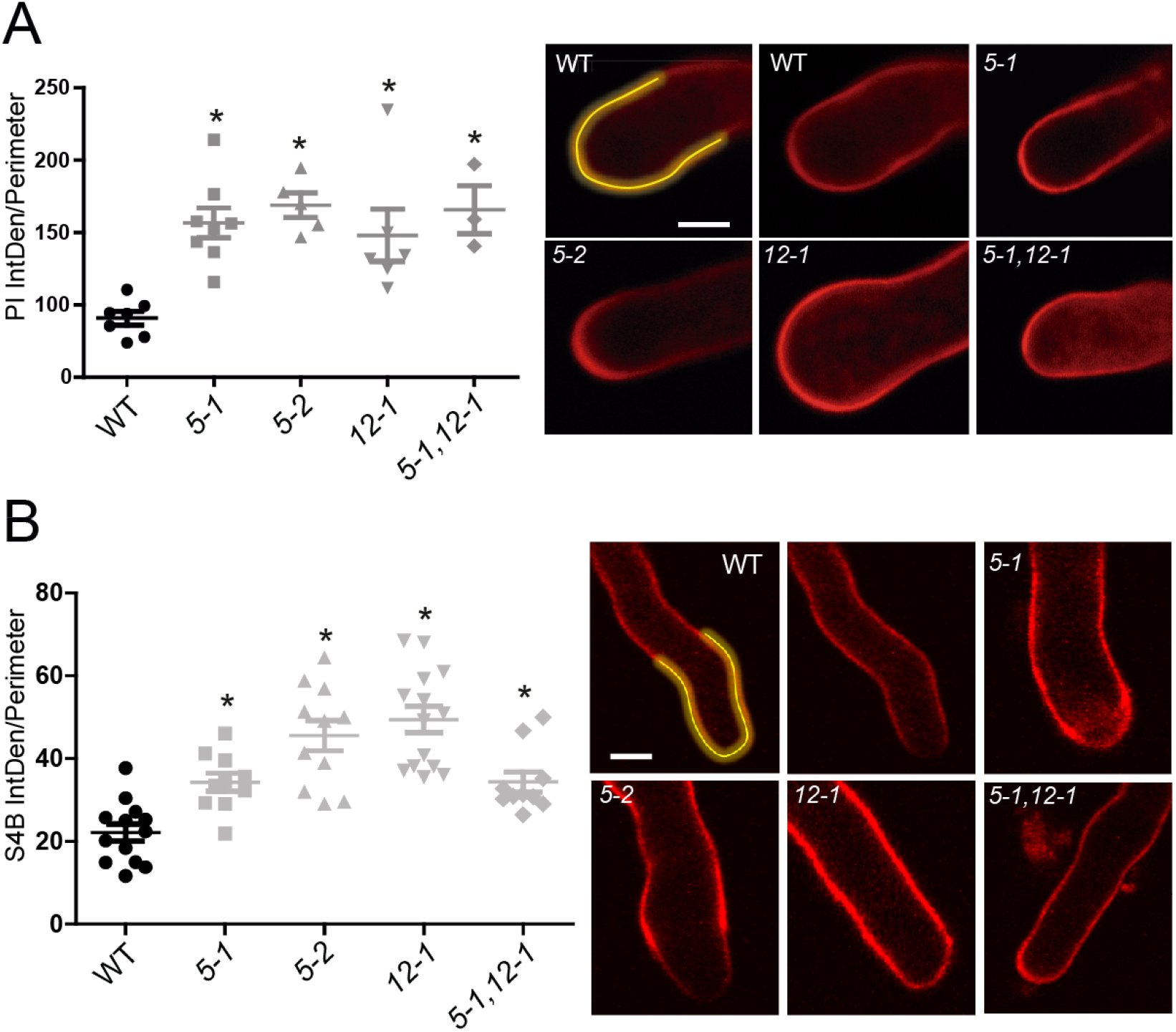
*Perk5, perk12 and perk5 perk12* mutants show greater deposition of pectin and cellulose in the cell wall of pollen tubes. (**A**) Determination of non-esterified homogalacturonans (pectins) by propidium iodide (PI) in the cell wall of pollen tubes germinated *in vitro*. The region outlined in yellow exemplifies the perimeter at which the fluorescence was measured for each pollen tube. Asterisks indicate significant differences from the WT according to one-way ANOVA test with p <0.05. On the right, representative images of *A. thaliana* pollen tubes stained with PI. Scale bar = 6 μm. **(B)** Determination of cellulose with Pontamine Fast Scarlet 4B (S4B) in the cell wall of pollen tubes germinated *in vitro*. The region outlined in yellow exemplifies the perimeter at which the fluorescence was measured for each pollen tube. Asterisks indicate significant differences from the WT according to one-way ANOVA test with p <0.05. On the right, representative images of *A. thaliana* pollen tubes stained with S4B. Scale bar = 10 μm.

ROS homeostasis is tightly controlled at multiple levels for a proper pollen tube growth (Potocky et al, 2007; Boisson-Dernier et al. 2009, 2013; Kaya et al., 2014; Lassig et al., 2014; Wudick and Feijó, 2014). Measurements of cytoplasmic ROS concentration in WT, and single mutant *perk5-1, perk5-2* and *perk12-1* and double mutant *perk5-1 perk12-1* pollen tubes was performed using 2,7-dichlorofluorescein diacetate probe (H_2_DCF-DA). Pollen tubes from the single mutant *perk5-1* and the double mutant *perk5-1 perk12-1* show significantly higher levels of ROS at the apical zone compared to WT pollen tubes, but not from *perk12-1* single mutant **(Figure 5A)**. However, because H_2_DCF-DA oxidation is irreversible and sensitive to different ROS and cannot be used to monitor ROS dynamics over time, we used HyPer, a genetically encoded YFP-based radiometric sensor, which is employed to detail the production of hydrogen peroxide (H_2_O_2_) in bacteria, animal and plant cells (Hernández-Barrera et al. 2015) including pollen (Mishina et al., 2013). Therefore, we generated stable mutant Arabidopsis *perk5-1, perk12-1* and *perk5-1 perk12-1* lines expressing HyPer under the control of the pollen specific *LAT52* promoter. **Figure 5B** shows that the average oscillating levels of H_2_O_2_ in growing pollen tubes of single *perk5-1* and *perk12-1*mutants and the double mutant *perk5-1 perk12-1* were higher compared to WT pollen. Taken together, these results suggest that PERK5 and PERK12 are part of the complex machinery that controls ROS homeostasis during pollen tube growth.

**Figure 5.**
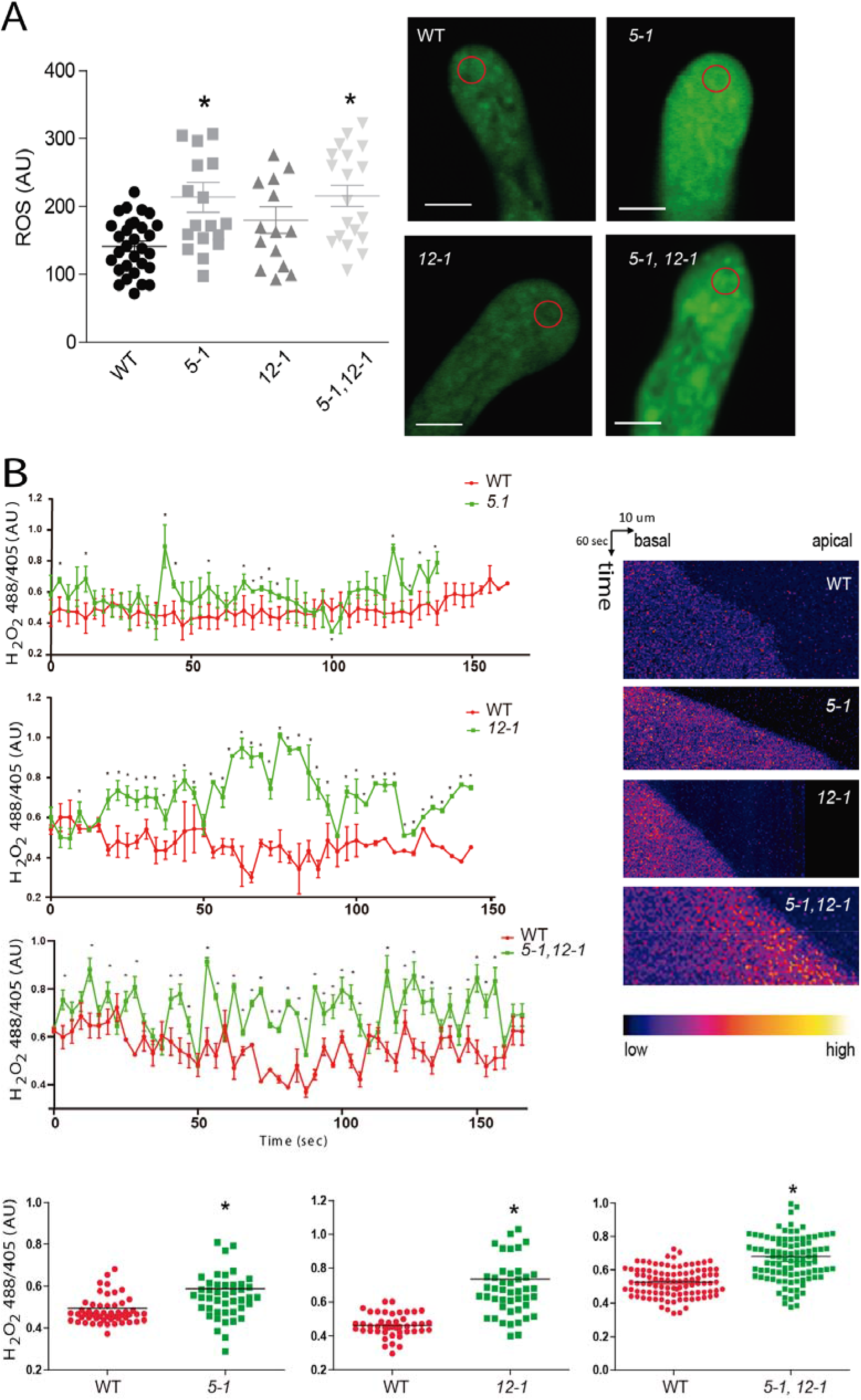
*Perk5, perk12 and perk5 perk12* mutants show higher levels of ROS at the apical zone of pollen tubes. (**A**) ROS levels in pollen tubes. Quantification of fluorescence intensity in pollen tubes stained with H_2_DCF-DA (n ≥ 15). Asterisks indicate significant differences with from the WT according to the one-way ANOVA test with p <0.05. On the left, representative images of pollen tubes stained with H_2_DCF-DA for 5 min. Red circles indicate the ROI where fluorescence was measured. Scale bar= 6 μm. AU= Arbitrary units. (**B**) Quantification of HyPer ratio (F488/F405) at the tip of growing WT, *perk5-1* and *perk12-1* pollen tubes. Data are shown as the mean± SEM of ratios over 150 sec. Asterisks indicate significant differences from the WT according to Student’s t test with p<0.05. On the right, kymographs representing the quantification. On the bottom, average signal was calculated based on each individual points.

## Discussion

Remodeling of cell wall structures, that is, degradation of old cell wall components, and biosynthesis, as well as incorporation of new material into the expanding cell wall, are necessary for proper pollen tube growth (Mollet et al, 2013; Hepler et al, 2013; Vogler et al, 2019). Monitoring the structure of the growing cell wall is also crucial during this process. Cell wall integrity (CWI) sensors perceive changes in cell wall structure and transduce this information into intracellular signaling cascades to adjust cell wall composition and, therefore, regulate polar growth. An elaborate CWI surveillance system consisting of apoplastic proteins and transmembrane receptors detects changes in cell wall homeostasis. The apical zone of growing pollen tubes is characterized by a gradient of cytoplasmic Ca^2+^ ions (_cyt_Ca^2+^) and ROS production. High levels of _cyt_Ca^2+^ in the tip zone trigger ROS production, in a reaction catalyzed by NADPH oxidases. Furthermore, high levels of ROS transiently elevate the concentration of _cyt_Ca^2+^ (Duan et al., 2014) by a still unknown mechanism. Respiratory Burst Oxidase Homolog protein H (*RBOHH*) and *RBOHJ* are proposed (Wu et al., 2010; Boisson-Dernier et al., 2013; Kaya et al., 2014; Lassig et al., 2014) to connect _apo_ROS production with the transient activation of plasma membrane Ca^2+^ channels (e.g. Cyclic Nucleotide Gated Channel 18 *[CNGC18]*, Glutamate-Like Receptor 1.2-3.7 [*GLR1.2* and *GLR3.7*] and Mechano-Sensitive Like channel 8 *[MSL8]*) in growing pollen tubes (Michard et al. 2011; Hamilton et al. 2015; Gao et al. 2016). In addition, *ANXUR1* (*ANX1*) and *ANXUR2 (ANX2)*, two RLKs of the pollen tube membranes, positively regulate *RBOHH* and *RBOHJ*, possibly through ROP signaling to produce oscillating ROS (Wudick and Feijó, 2014). Subsequently, ROS and the activate Ca^2+^-channels for calcium influx to fine-tune the tip-focused Ca^2+^ gradient, which in turn sustains exocytosis at the apical tip, enabling pollen tubes to elongate steadily without losing CWI. While ROS includes a variety of small molecules, H_2_O_2_ is the most stable and its production and transport need to be fine-tuned with high precision (Mangano et al. 2016). When *ANX1* and *ANX2* were absent or overexpressed, they have a direct impact on ROS production, suggesting a link between RLKs and RBOH activity in pollen tube growth (Boisson-Dernier et al. 2009, 2013). While ROS includes a variety of small molecules, H_2_O_2_ is the most stable and its production and transport need to be fine-tuned with high precision (Mangano et al. 2016). Abnormal levels of ROS, either lower or higher than under normal physiological conditions, inhibit or exacerbate pollen tube growth impacting on cell wall structure. Our results suggest that PERK5 and PERK12 are involved in the control of ROS levels during pollen tube growth. Likewise, *perk13* mutant (*rsh10*) also showed elevated ROS levels in roots (Hwang et al. 2016) suggesting that *PERK13* represses *RBOHC* activity in root hair cells (Foreman et al. 2003). There may be a more general link between PERKs and RBOHs, although their association during pollen tube growth remains to be established. In the recent years, great advances on the CWI pathway have been accomplished. CWI sensing involves several receptor kinases at the plasma membrane as well as apoplastic proteins that can bind cell wall components and trigger directly or indirectly intracellular processes. Among these components, WAKs and CrRLK1L as receptors while peptides of the Rapid Alkalinization Factors (RALFs) and proteins of the Leucine-rich repeat extensins (LRXs) families, as apoplastic proteins have been characterized (Xu et al. 2008; Kohorn 2016; Nissen et al. 2016; Franck et al. 2018; Mecchia et al. 2017; Sede et al. 2018; Wang et al. 2019). In this complex scenario, we have identified two surface RLKs (PERKs) that could contribute to the ROS-regulated polar growth of pollen tubes which could be good candidates for CWI sensors at the pollen tube tip.

Here, we showed that *PERK5* and *PERK12*, two Arabidopsis pollen expressed PERKs, are necessary for pollen tube growth. We found that *perk5-1 perk12-1* double mutant displayed shorter pollen tubes and showed a reduced number of seeds per silique. These results could be explained by the observation of an accumulation of pectins, in particular, non-methylated pectins, in the apical and subapical zone of mutant pollen tubes. This low level of esterification would allow pectins to cross-link with Ca^2+^, giving rigidity to the cell wall, which would explain the shorter pollen tube phenotype of the double mutant *perk5-1 perk12-1*. According to this, the increase in cellulose abundance at the pollen tube apical zone of the *perk* mutant plants would also explain the observed phenotypes since a high cellulose content is associated with a decrease in the pollen tube growth rate (Mollet et al., 2013). Our findings provide new insight into the function of PERK proteins, possibly as part of the CWI sensor pathway.

Very little is known about downstream components of PERKs proteins. *KIPK1* and *KIPK2*, which belong to the AGC1 subgroup of AGCVIII family (Zegzouti et al. 2006; Rademacher and Offringa, 2012), were previously identified as interactors by their N-terminal domains to the cytosolic kinase domains of PERK8, PERK9 and PERK10 (Humphrey *et al*., 2015; Li et al. 2017). Since AGCVIII plant proteins are most closely related to animal PKA and PKC, involved in regulation of polar growth, it is not surprising that the double mutant for the *AGC1.5* and *AGC1.7* pollen genes exhibit defective pollen tube growth (Zhang et al. 2009). The fact that the physical interaction between AGC1 kinases and PERKs is maintained as a functional unit in different plant cell types, makes pollen AGC1.5 and AGC1.7 good candidates as interactors with PERK5 and PERK12, and together would be involved in the same signaling pathway that controls pollen tube growth. Further characterization of PERKs regarding their subcellular localization in pollen tubes, a higher order of multiple PERK mutants and their association to AGC1 proteins, will be the best approach to understand in detail the role of pollen PERKs during pollination.

## Conclusions

We identify *PERK5* and *PERK12*, two *A. thaliana* pollen expressed PERKs, as necessary for pollen tube growth. We found that *perk5-1 perk12-1* double mutant displayed shorter pollen tubes, a deficient pollen transmission and a reduced number of seeds per silique. In addition, these mutants showed changes in their cell wall composition and disturbed ROS homeostasis that directly affect the pollen tube expansion rate. Our findings provide new insights into the function of PERK proteins, possibly as part of the CWI sensor pathway.

## Methods

### Plant material and growth conditions

*A. thaliana* seeds were germinated in 0.5X MS culture medium (Murashige and Skoog 1962) containing 1% agar in an incubator at 22°C under long day conditions (16h light/8h dark). 10-day-old seedlings were transferred to soil and grown under the same conditions as described above.

### Identification of PERK T-DNA insertional lines

T-DNA insertional lines for each PERK gene were obtained from ABRC (Arabidopsis Biological Resource Center). For identification of T-DNA knock-out lines, genomic DNA was extracted from rosette leaves. Single and multiple T-DNA insertions in the target genes were confirmed by PCR. Homozygous lines were isolated for the genes included in this study. *Arabidopsis thaliana* Columbia-0 (Col-0) was used as the WT genotype in all experiments. Mutant lines and the primers used for genotyping the T-DNA lines are listed in **Table S1**.

### RT-PCR analysis

Pollen from mature flowers of WT and *perk5-1, perk5-2, perk12-1*, mutants were germinated *in vitro* and total RNA from emerging pollen tubes were extracted with the RNAzol method (MRC, Inc) according to the manufacturer’s instructions. For cDNA synthesis, 100 ng of the recovered RNA was used as a template for M-MLV reverse transcriptase (Promega). The PCR reactions were performed in a T-ADVANCED S96G (Biometra) using the following amplification program: 4 min at 95°C, followed by 35 cycles of 30 sec at 95°C, 30 sec at 57°C and 30 sec at 72 °C. *PP2A* served as an internal standard. All the primers used are listed in **Table S1**.

### *In vitro* and *semi-in vivo* pollen germination

To assay *in vitro* pollen germination, pollen was collected and cultured in a medium containing 10% sucrose, 0.01% H_3_BO_3_,1 mM MgSO_4_, 5 mM CaCl_2_, 5 mM KCl and the pH was adjusted to 7.5 (Boavida & McCormick., 2007). For semi-solid medium 1% low-melting agarose was added. Pollen grains were germinated at 22°C with 100% relative humidity in an incubation chamber. For quantitative analysis of pollen tube length in T-DNA mutants and WT, 200 pollen tubes were measured (number of plants = 11) from mature flowers. Images were taken using a Zeiss microscope Axio Imager A2 (Carl Zeiss). Values are reported as the mean ±SEM using the Image J 1.47d software. For kinetics analysis pollen tubes were imaged and measured every 10 min and growth rates were calculated. For pollen germination experiments *in vivo*, pre-emasculated mature flowers were pollinated either with WT or mutant pollen. After 3 h and 12 h pistils were isolated and fixed with a mixture of acetic acid: ethanol (3:1), rehydrated with an ethanol series (ethanol 70%, 50%, 30%) cleared with 8 M sodium hydroxide and stained with decolorized aniline blue (Mori *et al*., 2006). Images of stained pistils were taken with a Zeiss microscope Axio Imager A2 under UV light.

### ROS measurements with H_2_DCF-DA

After 3 h of *in vitro* germination on semi-solid medium, pollen tubes were incubated for 5 min with 50 μM H_2_DCFDA (Molecular Probes, Invitrogen, C6827) at room temperature, and then were washed away with fresh dye-free medium before imaging. Pollen tubes stained with H_2_DCF-DA were imaged with a Zeiss Meta 510 LSM confocal microscope. For H_2_DCF-DA stained pollen tubes, a circular ROI away from the tip was chosen to measure apical cytosol intensity. All dye-derived fluorescence intensities were measured using the ImageJ 1.47d software after background subtraction. Pollen tubes of different genotypes were all imaged and quantified under the same conditions.

### H_2_O_2_ imaging with HyPer sensor

Fluorescence in growing pollen tubes of WT and T-DNA lines expressing HyPer were acquired with Zeiss Meta 510 LSM confocal microscope (Carl Zeiss) and were quantified (ImageJ 1.47d software) in the same conditions. HyPer fluorescence was acquired with the sequential mode: excitation at 488 nm and emission between 500–540 nm for F488 and excitation at 405 nm and emission between 500–540 nm for F405. A circular ROI at the apex was drawn for measurement of apical cytosol intensity for each single time point of each pollen tube. All ratiometric measurements (F488/F405) were determined with ImageJ 1.47d software and its Ratio Plus Manager plugin after background subtraction. Kymograph pictures were generated with the Multiple Kymograph plugin.

### GUS assay

The regulatory region (2.0 kb) of *PERK* was cloned into a pENTRY-D-TOPO, and then was subcloned into pMDC163. Plants were transformed and transgenic plants were selected. To visualize the activity of p*PERK5::GUS* reporter, inflorescences and seedlings of transgenic plants were subjected to GUS staining, according to Donnelly *et al*. (1999). Inflorescences were cleared with chloral hydrate clearing solution (8 g of chloral hydrate, 1 ml of glycerol, and 2 ml of H_2_O) for 30 min at room temperature before imaging. Bright-field images were taken with an MVX10 Research Macro Zoom Microscope (Olympus).

## Acknowledgements

We would like to acknowledge L. Cárdenas for providing LAT52::HyPer sensor line. We thank ABRC (Ohio State University) for providing T-DNA seed lines. J.M.E. and J.P.M are investigators of the National Research Council (CONICET) from Argentina. This work was supported by a grants from the Agencia Nacional de Promoción Científica y Tecnológica (ANPCyT) (PICT2016-0132, PICT2017-0066), Fondo Nacional de Desarrollo Científico y Tecnológico (1200010), and Instituto Milenio iBio – Iniciativa Científica Milenio, MINECON to JME, and from the ANPCyT (PICT2017-0076, PICT2018-0504) to JPM. The authors declare that they have no competing financial interests. Correspondence and requests for materials should be addressed to JPM or JME.

## Author’s contribution

C.B. and A.S. performed all the experiments, analyzed the data and wrote the paper. M.A.M, S.M., E.M., S.P.D.J., S.M.V., J.D.S.S. analyzed the data. J.P.M. and J.M.E. designed research, analyzed the data, supervised the project, and wrote the paper. All authors commented on the results and the manuscript. This manuscript has not been published.

---

**Supplementary Figure 1.**
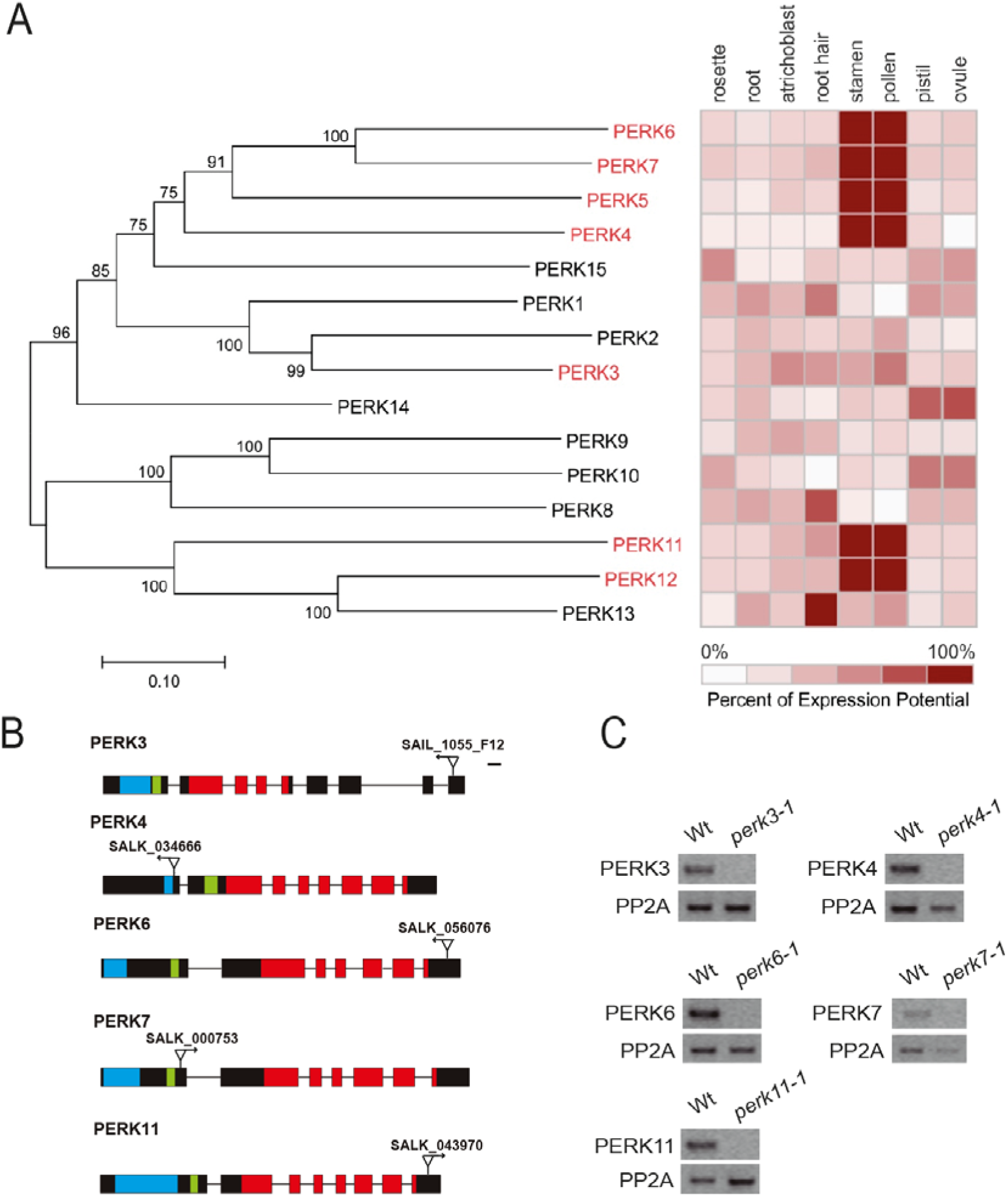
PERKs expression patterns and characterization. **(A)** Phylogenetic tree of Arabidopsis PERK proteins (left) combined with relative expression (right). The phylogenetic analysis was carried out with MEGA6 (Tamura et al., 2007) using the Maximum Similarity method (Maximum Likelihood) (Saitou and Nei, 1987). The numbers in the nodes indicate the bootstrap values obtained for 1000 iterations. Scale represents the evolutionary distance, expressed as the number of substitutions per amino acid. Relative expression of *PERKs* in Arabidopsis in different tissues is shown. **(B)** Schematic representation of PERK genes showing the extracellular (EXT)-domains with Ser-Pro_(3–5)_ repeats (light blue), transmembrane (green) and kinase domains (red) coding sequences. Introns (thin lines), exons (rectangles), and positions of T-DNA insertions are also indicated. Bars = 100 bp. **(C)** RT-PCR of *perk* mutant lines. RNA was extracted from pollen tubes germinated *in vitro. PP2A* was used as control.

**Supplementary Figure 2.**
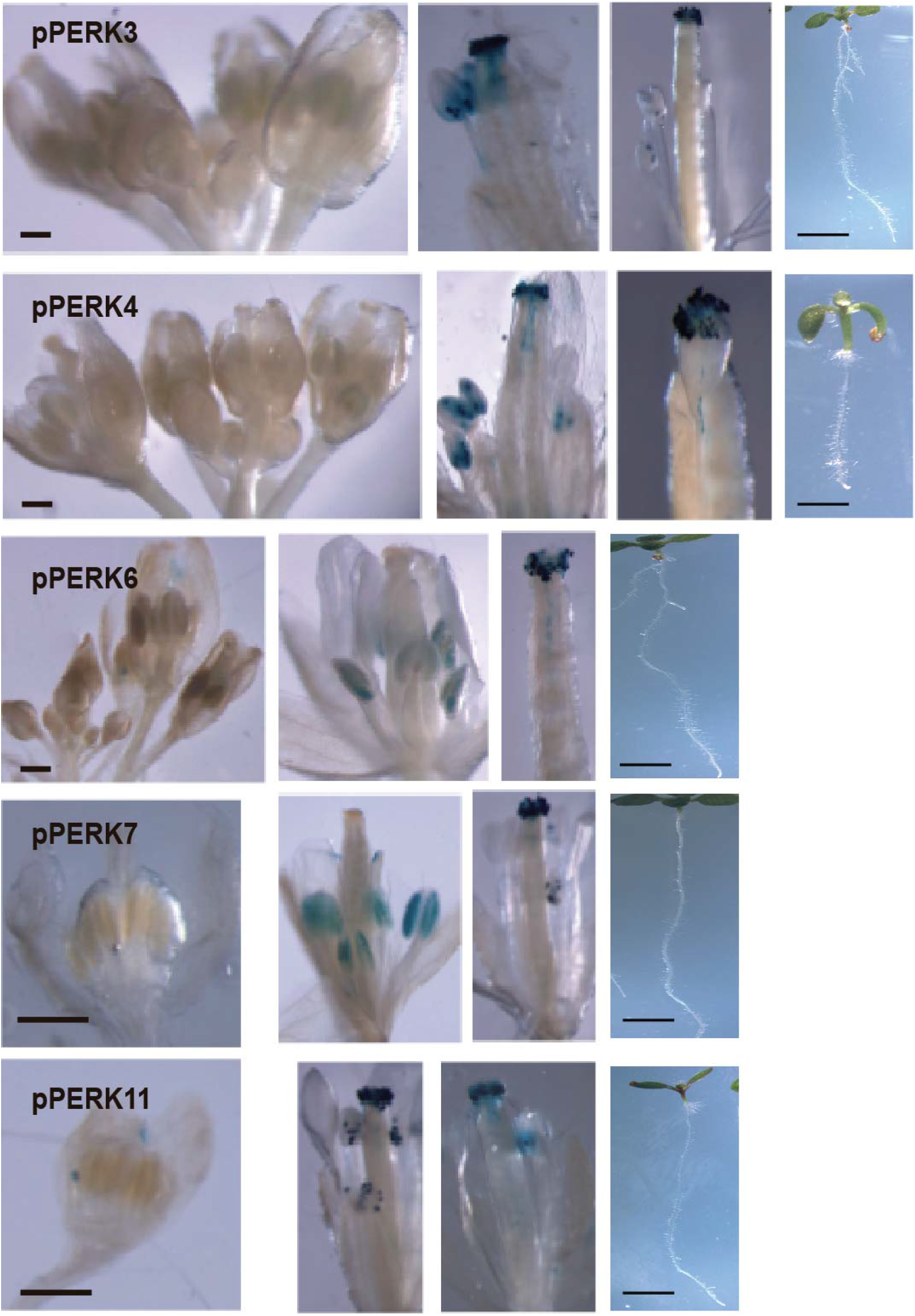
Expression pattern of pPERK::GUS reporters. GUS activity in the *pPERK::GUS* plant lines. GUS activity in plants for *pPERK3::GUS, pPERK4::GUS, pPERK6::GUS, pPERK7::GUS and pPERKll::GUS* are shown. GUS activity was visualized in mature pollen grains and in pollen tubes growing through the style. Scale = 100 μm. Right panels visualize seedlings of each transgenic line in which GUS activity was not detected. Scale bar: 2mm.

**Supplementary Figure 3.**
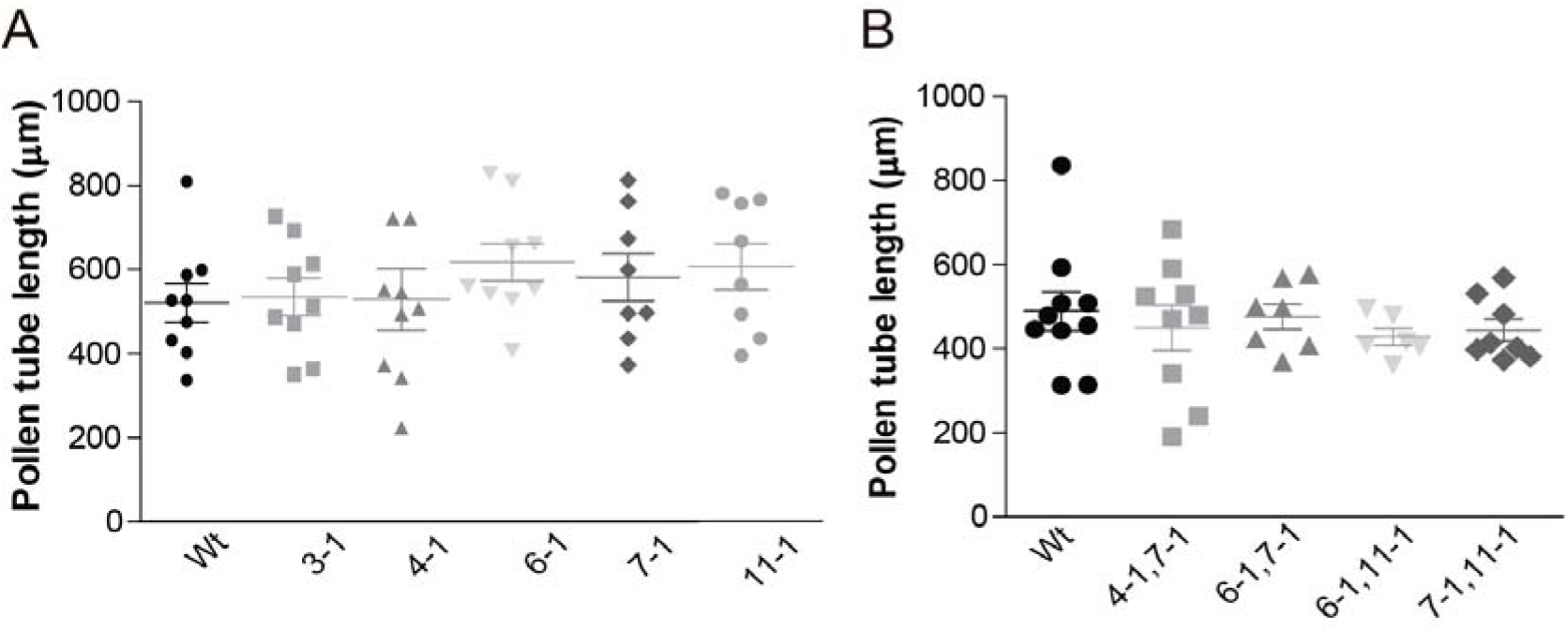
Analysis of pollen germination and pollen tube length in single and double *perk* mutants. **(A)** Quantification of pollen tube length in single mutants after 3h of *in vitro* germination. Data are shown as mean ± SEM (n ≥ 10). **(B)** Quantification of pollen tube length in double mutants after 3h of *in vitro* germination. Data are shown as mean ± SEM (n ≥ 8). No significant differences were detected in all the mutants (for **A-B)** when compared to the WT according to ANOVA test with p <0.05.

**Supplementary Table 1.**
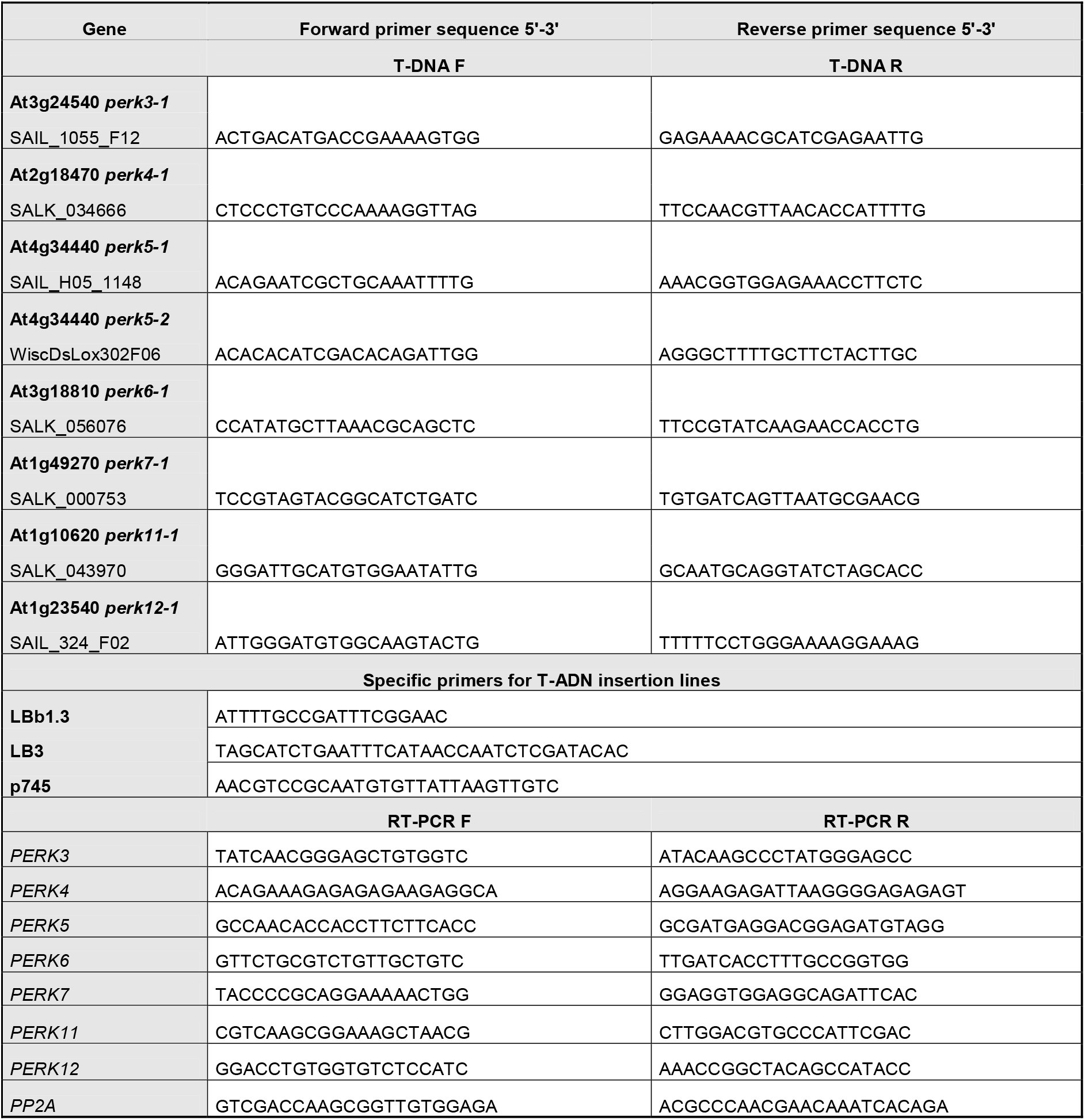
List of primers used in this study.

